# The translation attenuating arginine-rich sequence in the extended signal peptide of the protein-tyrosine phosphatase PTPRJ/DEP1 is conserved in mammals

**DOI:** 10.1101/2020.09.07.285544

**Authors:** Luchezar Karagyozov, Petar Grozdanov, Frank-D. Böhmer

**Affiliations:** Institute of Molecular Cell Biology, CMB, Jena University Hospital, Jena, Germany; Department of Cell Biology & Biochemistry, School of Medicine, Texas Tech University Health Sciences Center, Lubbock, USA

## Abstract

The signal peptides, present at the N-terminus of many proteins, guide the proteins into cell membranes. In some proteins, the signal peptide contains an extended N-terminal region and a recessed hydrophobic signal sequence. Previously, it was demonstrated that the N-terminally extended signal peptide of the human *PTPRJ* contains a cluster of arginine residues, which attenuates translation. The analysis of the orthologous sequences revealed that this sequence is highly conserved among mammals. The *PTPRJ* transcripts in placentals, marsupials, and monotremes encode a stretch of 10 – 14 arginine residues, positioned 11-12 codons downstream of the initiating AUG. The remarkable conservation of the repeated arginine residues in the *PTPRJ* signal peptides points to their key role. Further, the presence of an arginine cluster in the extended signal peptides of other proteins (E3 ubiquitin-protein ligase, NOTCH3) is noted and indicates a more general importance of this cis-acting mechanism of translational suppression.

## Introduction

After the start of translation, translation inhibition due to the interaction between the nascent polypeptide chain and the ribosome is reported (Ito and Chiba, 2013; Wilson et al., 2016). In eukaryotes, transient elongation arrest may be caused by consecutive prolines (Artieri and Fraser, 2014) or by an array of positively charged amino acids (Lu and Deutsch, 2008). In a limited number of proteins, the signal peptides are longer than the canonical 20 −25 residues; they consist of more than forty amino acid residues and contain an N-terminal extension and a recessed hydrophobic signal sequence (Hiss et al., 2008). This extension provides a convenient space for positioning of a translation attenuating amino acid sequence.

The human receptor-like protein tyrosine phosphatase, type J (PTPRJ, PTPReta, or DEP1) provides an example of the presence of a down-regulating sequence within the signal peptide. The human PTPRJ is a receptor-like protein tyrosine phosphatase of the R3 subtype, characterized by an extracellular region, containing several tandem fibronectin type III (FNIII) domains, a single transmembrane region, and a single cytoplasmic catalytic domain (Andersen et al., 2001). In human embryonic lung fibroblasts, the *PTPRJ* expression and activity were dramatically increased when cells approached confluence in comparison to sparse cells, suggesting a possible role in cell-density-dependent inhibition of proliferation. Thus, the name high cell density-enhanced phosphatase-1 or DEP1 was proposed (Ostman et al., 1994). *PTPRJ* is expressed in a variety of normal tissues, notably in hematopoietic cells (CD148 antigen), in epithelial tissues, including those of the digestive tract, in the vascular endothelium (Autschbach et al., 1999). Data suggest that *PTPRJ* is a tumor suppressor in different tissues. In mice, the gene encoding PTPRJ was mapped to a colon cancer susceptibility locus (Scc1) (Ruivenkamp et al., 2002). Negative regulation of the signaling of several receptor-tyrosine kinases (RTKs), including the epidermal growth factor receptor (EGFR) (Tarcic et al., 2009), the platelet-derived growth factor receptor (Petermann et al., 2011), and Fms-like tyrosine kinase 3 (Arora et al., 2011) may be important in this context. A metabolic function of PTPRJ is indicated by the negative regulation of insulin receptor and leptin receptor signaling (Shintani et al., 2015, Krüger et al., 2015, Shintani et al., 2017). PTPRJ was also identified as an effective activator of Src-kinase in different cell types. This function of PTPRJ is important for platelet activation and thrombosis (Senis et al., 2009, Nagy et al., 2020), for efficient angiogenesis (Fournier et al., 2016), and for regulating airway hyper-responsiveness (Katsumoto et al., 2013).

Previous experiments (Karagyozov et al., 2008) showed that: (1) the human *PTPRJ* transcripts predominantly initiate translation at the first AUG in a favorable context, numbered AUG_190_. This results in a PTPRJ pre-protein with an N-terminally extended signal peptide and a recessed hydrophobic signal sequence. (2) The N-terminal extension contains an unusual arginine-rich cluster (RRTGWRRRRRRRR); its translation inhibits the overall PTPRJ expression.

To elucidate the importance of these findings it was of interest to examine the *PTPRJ* transcripts in mammals for sequences encoding the attenuating arginine-rich cluster. In the present paper, *PTPRJ* orthologs from placental mammals, marsupials, and monotremes were compared. The results revealed a similarity in the architecture of the *PTPRJ* transcripts. Several conserved features were noted: (1) uORFs are not present in the transcripts; (2) the *PTPRJ* transcripts encode signal peptides with N-terminal extension and a recessed hydrophobic signal sequence; (3) the extended signal peptides contain the attenuating arginine cluster. The remarkable evolutionary conservation of the attenuating sequence emphasizes the importance of suppressing the PTPRJ translation by the nascent peptide chain.

## Materials and methods

### Orthologous genes and data evaluation

The NCBI gene database was used to search for *PTPRJ* orthologs in mammals. The 5’ end regions of the *PTPRJ* transcripts were examined for the presence of initiator AUG codons to determine the N-terminal end of the encoded proteins. The presence of a signal hydrophobic amino acid sequence and the signal-peptidase cleavage sites were determined *in silico* using the SignalP 5.0 web server at https://services.healthtech.dtu.dk/service.php?SignalP-5.0 (Armenteros et al., 2019). In the SignalIP algorithm, the input sequence has an upper limit of 70 residues, thus, initially, the N-terminal amino acids were examined, and then - the adjoining downstream region. The distribution of the elongating and initiating ribosomes in transcripts encoded by exon 1 (450 bp) of the human *PTPRJ* on chromosome 11 was visualized by the genome browser https://gwips.ucc.ie/ (Michel et al., 2014). BLAST, PHI-BLAST (NCBI) and Clone Manager Suite 8 (Scientific and Educational Software) were used to search the database, compare and analyze the nucleotide and amino acid sequences.

### Protein-tyrosine phosphatase PTPRJ/DEP1 sequences

The analyzed 5’ end regions of the *PTPRJ* transcripts were from ten species: five placental mammals, four marsupials, and one monotremes.

### Placentals

Primates - *Homo sapiens* (human), mRNA transcript variant 1: NM_002843.4, protein: NP_002834.3

Rodents - *Mus musculus* (house mouse), mRNA transcript variant 1: NM_008982.6, protein: NP_033008.4

Cetaceans - *Delphinapterus leucas* (beluga whale), mRNA: XM_030764039.1, protein: XP_030619899.1

Ruminants – *Bos taurus* (cattle), mRNA: XM_024975918.1, protein: XP_024831686.1

Carnivore - *Enhydra lutris* (sea otter), mRNA: XM_022506988.1, protein: XP_022362696.1

### Marsupials

*Monodelphis domestica* (gray short-tailed opossum) - mRNA: XM_016422845.1, protein: XP_016278331.1

*Phascolarctos cinereus* (koala) - mRNA transcript variant X1: XM_021009664.1, protein: XP_020865323.1

*Sarcophilus harrisii* (Tasmanian devil) - mRNA transcript variant X1: XM_031941343.1, protein: XP_031797203.1

*Vombatus ursinus* (common wombat) - mRNA transcript variant X1: XM_027837689.1, protein: XP_027693490.1

### Monotremes

*Ornithorhynchus anatinus* (platypus) – mRNA transcript variant X2: XM_029060042.1, protein: XP_028915875.1

### E3 ubiquitin-protein ligase (ZNRF3) sequences

*Homo sapiens* (human), mRNA transcript variant 1: NM_001206998.2, protein: NP_001193927.1

*Mus musculus* (house mouse), mRNA transcript variant 1: NM_001080924.2, protein: NP_001074393.1

### Notch receptor 3 (NOTCH3) sequences

*Homo sapiens* (human), mRNA transcript: NM_000435.3, protein: NP_000426.2 *Mus musculus* (house mouse), mRNA: NM_008716.3, protein: NP_032742.1

## Results and discussion

### The *PTPRJ* orthologs in mammals

The NCBI gene database lists 123 mammalian orthologs of the human *PTPRJ:* placental mammals - 118 orthologs, marsupials - 4, monotremes - 1. We analyzed five orthologs from major groups of placentals and all orthologs from marsupials and monotremes.

In some mammalian species, different splice variants encoding different PTPRJ isoforms are listed. We examined all isoforms for the presence of a hydrophobic signal sequence near the N-terminus. Only isoforms with a predicted cleavable signal peptide were analyzed further.

### The AUG initiator codons in mammalian *PTPRJ* transcripts

The arrangement of the AUG triplets at the 5’ end of the mammalian *PTPRJ* mRNA is shown in Fig. 1. In all transcripts, the AUG codons are in-frame with the mature protein, with no intervening stop codons between them (see also Suppl. Fig. 1, 3 and 5). Thus, no uORFs exist in the mammalian *PTPRJ* transcripts.

**Figure 1.**
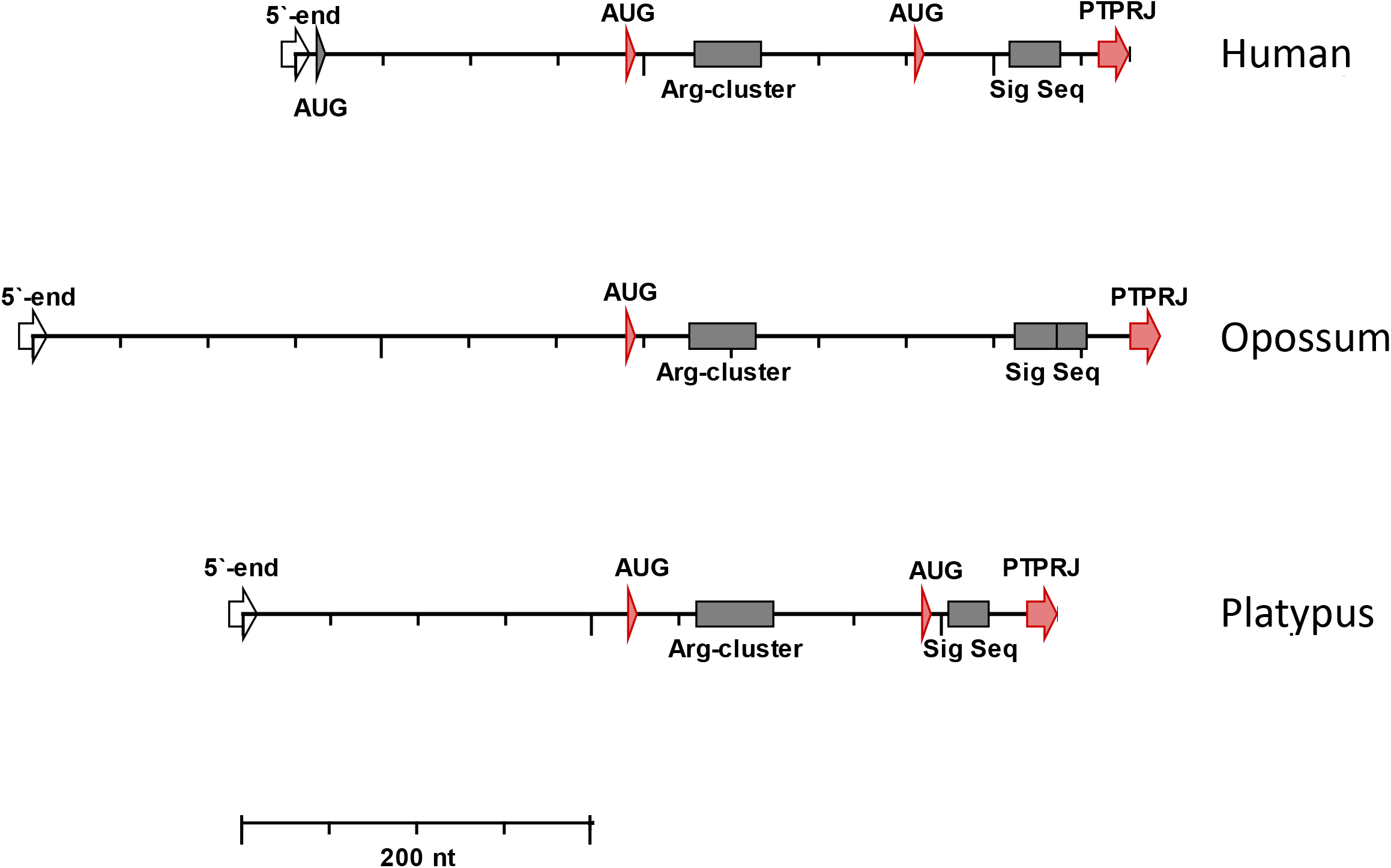
Sequence pattern of the 5’-end of the mammalian *PTPRJ* transcripts. The AUG codons are indicated. The position of the transcripts is adjusted to align the first initiating AUGs. Arg-cluster – repeated Arg residues; Sig Seq – region of nonpolar amino acids; PTPRJ – the mature protein

The number of the initiating AUGs differs between mammalian subdivisions; the placentals have three AUGs, the monotremes – two, and the marsupials – only one. Remarkably, this AUG is homologous to the previously identified (Karagyozov 2008) preferred starting site for translation in humans AUG_190_ (see below).

It is firmly established that the nucleotides surrounding the AUG codons strongly affect initiation efficiency. Mutagenesis experiments with transfected COS cells (Kozak, 1986) established an optimal context sequence for initiation (RCC<u>**AUG**</u>G, R is A or G). The nucleotides at positions <u>-3</u> and <u>+4</u> (the A in the AUG codon is <u>+1</u>) are of particular importance. The AUG context is categorized as strong (both crucial positions match the optimal sequence), favorable (only one match), or weak (no match at positions <u>-3</u> and <u>+4</u>).

In mammalian *PTPRJ,* the nucleotide context of the AUG codons varies according to their scanning order (Table 1). In placentals, the AUG close to the 5’ cap (6 – 15 nucleotides) is scanned first; it is in a weak context (CGC<u>**AUG**</u>A). The next AUG is in a favorable context with G at <u>-3</u> and U at <u>+4</u> (GCC<u>**AUG**</u>U); the context of the third AUG is also favorable, however with A at <u>+4</u> (GCC<u>**AUG**</u>A). In monotremes, the context of the AUGs is favorable (GCC<u>AUG</u>U and GCC<u>AUG</u>A, respectively). The marsupials possess a single AUG, which is in a favorable context (GCC<u>**AUG**</u>U).

**Table 1.** Context of the AUG codons in the 5’ end of the *PTPRJ* transcripts. (The AUG codons are arranged in 5’ to 3’ direction according to the movement of the scanning complexes).

| Organism | 1<sup>st</sup> AUG | 2<sup>nd</sup> AUG | 3<sup>rd</sup> AUG |
| --- | --- | --- | --- |
| Consensus | CGC<u>AUG</u>A | GCC<u>AUG</u>U | GGC<u>AUG</u>A |

Monotremes.
| Organism | 1<sup>st</sup> AUG | 2<sup>nd</sup> AUG |
| --- | --- | --- |
| Platypus | GCC<u>AUG</u>U | GGC<u>AUG</u>A |

Marsupials.
| Organism | 1<sup>st</sup> AUG |
| --- | --- |
| Consensus | GCC<u>AUG</u>U |

To summarize, in all mammals, the AUG in a favorable context, at which the scanning 40S subunits arrive first, is with U in position <u>+4</u>. In placentals, the transcripts contain a preceding AUG in a weak context. In monotremes and placentals, the transcripts possess a downstream AUG in a favorable context, but with A in position <u>+4</u>. The functional significance of these differences is unknown. One may speculate that they reflect subtle differences in expression regulation.

### The efficiency of the AUG initiator codons in the human *PTPRJ* transcripts

In humans, there are three in-frame AUG codons positioned upstream of the hydrophobic signal sequence (Fig. 1). In previous experiments, the efficiency of each of the human initiator codons was tested in reporter constructs (Karagyozov et al., 2008). Briefly, the outcomes were (1) when all three AUGs were mutated no reporter activity was detected; (2) the efficiency of the first initiator (AUG_13_) was the weakest, and (3) the efficiency of the next two initiators (AUG_190_ and AUG_355_) appeared similar. Translation of the *PTPRJ* mRNA started predominantly at the first initiator in a favorable context (AUG_190_).

These results are in agreement with published RiboSeq data (Fig. 2). The sequences encoded by the first exon of the human *PTPRJ* do not support appreciable non-canonical translation initiation. The first initiator (AUG_13_), which is in a weak context, is not “tight”; it leaks scanning complexes downstream towards the second initiator. According to the profiles of the initiating ribosomes, the third initiator (AUG_355_) is not active.

**Figure 2.**
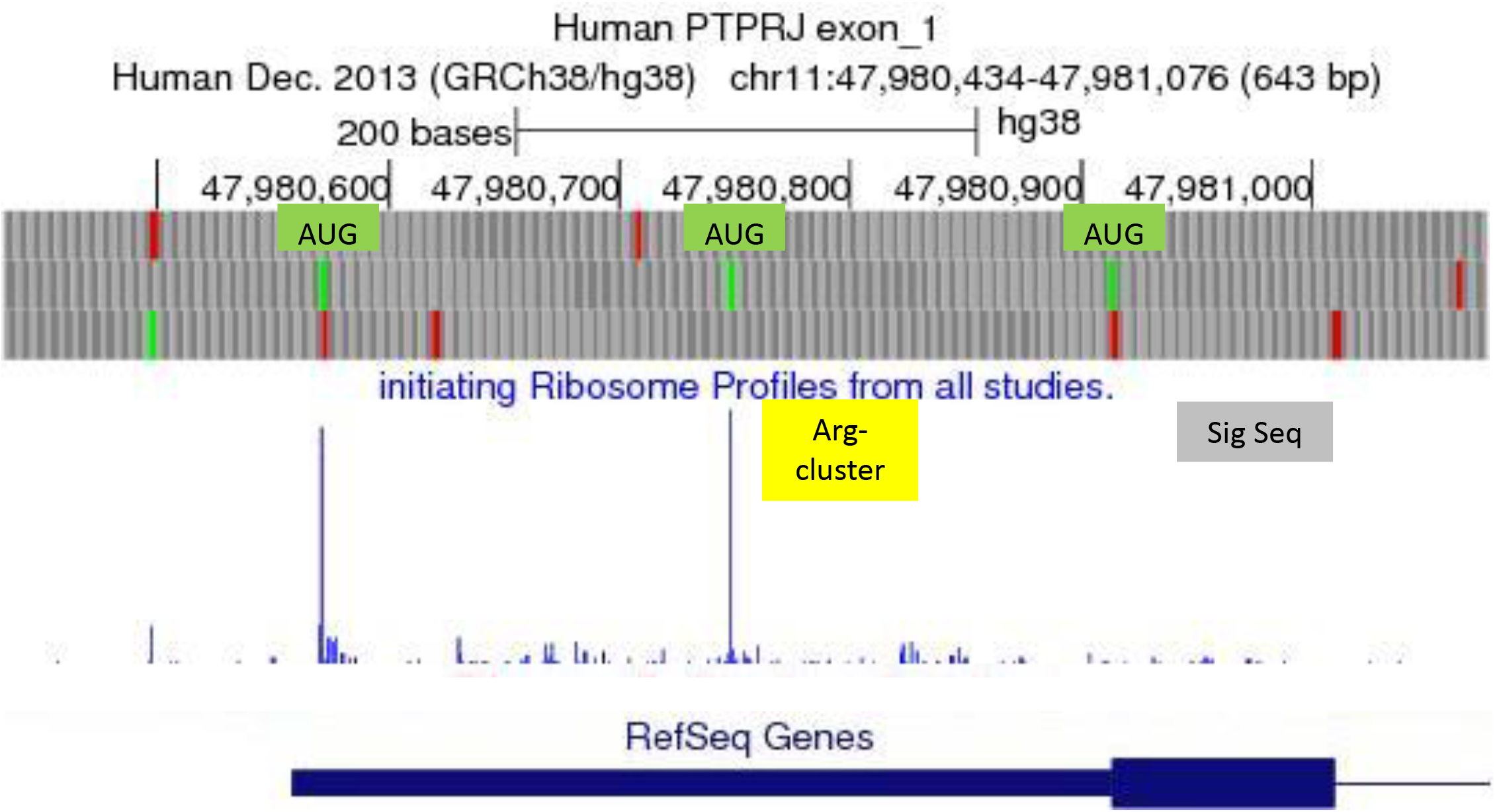
Ribosome profiling of *hPTPRJ,* exon 1. Top – the AUGs in reading frame 2 are shown (green). Middle - RiboSeq data, initiating ribosomes. The positions of the Arg-cluster (yellow) and the signal sequence (grey) are indicated. Bottom – the RelSeq gene, exon 1.

### The mammalian *PTPRJ* transcripts code for signal peptides with N-terminal extension

The N-termini of the proteins targeted for secretion or membrane integration, harbor a short amino acid sequence – the signal peptide – instrumental for the translocation of the proteins into the ER membranes. In most cases, the signal peptide is 20 – 25 amino acids long. It contains a positively charged N-terminal region (1 – 5 residues), a hydrophobic signal sequence (7 - 15 residues), and a signal-peptidase recognition site (von Heijne, 1990).

In several proteins, however, the signal peptide is N-terminally extended and the hydrophobic signal sequence is recessed (Hiss et al., 2008). Recent experiments showed that the N-terminal extension of the signal peptides is irrelevant for the signal sequence function (Meyer et al., 2018).

The signal peptides of the mammalian *PTPRJ* proteins are with an N-terminal extension (Fig. 1). In marsupials, the signal peptides are 94-103 residues long (Suppl. Fig. 4). In the platypus (monotremes), the *PTPRJ* mRNA encodes two possible signal peptides. The translation launched from the first AUG results in an N-terminally extended signal peptide of 76 amino acids (Suppl. Fig. 6). In placentals, the AUG initiating codons are three. According to the profiles of the initiating ribosomes, the third initiator (AUG_355_) is not active (Fig. 2). The first and second initiators produce N-terminally extended signal peptides composed of 147 - 150 and 87 – 92 amino acid residues, respectively (Suppl. Fig. 1 and 2).

### The arginine-rich sequence in the N-terminally extended signal peptides is highly conserved

The human *PTPRJ* transcripts encode an extended signal peptide, which contains repeated arginine residues (RRTGWRRRRRRRR). Earlier it was demonstrated that the arginine cluster attenuates translation (Karagyozov et al., 2008). A comparison between mammalian orthologs revealed the presence of a similar sequence in the extended signal peptides of all mammals (Fig. 3). Differences are minor. In placentals, the sequence of arginine residues is with three intervening amino acids (TGW/G). In marsupials, the string of arginine residues is interrupted by two amino acids (S/TW). In platypus, the arginine-rich cluster is 15 amino acids with one interruption (G). The arginine repeats in mammals are positioned 11-12 residues downstream of the initiating methionine.

**Figure 3.**
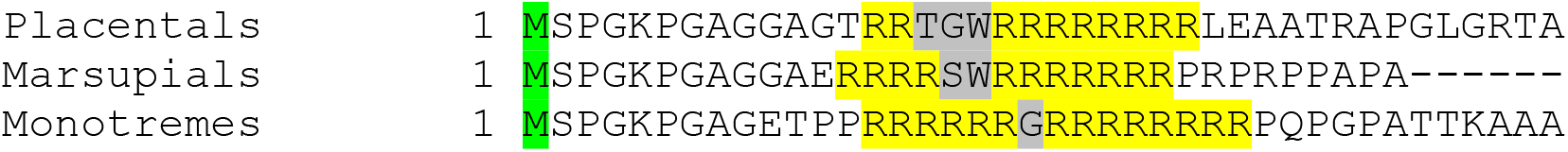
Comparison of the arginine cluster in the N-terminally extended signal peptides in mammals (placentals and marsupials - consensus sequences, monotremes – platypus, see Suppl. Fig. 2, 4 and 6). The initiating Met residues (green), the conserved Arg-residues (yellow) and the intervening amino acids (grey) are marked.

The conservation of the composition and location of the arginine-rich cluster emphasizes the functional importance of these features. In the absence of uORFs, this seems to be a necessary mechanism to down-regulate PTPRJ expression.

### The mechanism of translation attenuation and potential biological significance

Previous experiments showed that: (1) the translation attenuation is not due to the presence of rare codons; (2) elimination of the repeated arginine residues by frame-shift mutations (plus and minus) is sufficient to up modulate *PTPRJ* expression (Karagyozov et al., 2008).

Most likely, the inhibition of expression is due to the positive charge of the arginine residues. Stalling of ribosomes at positively charged residues, due to electrostatic interactions with the negatively charged exit tunnel was described in model experiments (Lu and Deutsch, 2008). More recently RiboSeq data were interpreted to show that ribosomes in yeast and mammals stall at positively charged amino acids (Charneski and Hurst, 2013: Sabi and Tuller, 2015). The ribosome exit tunnel accommodates 30 – 40 amino acid residues (Ito and Chiba, 2013; Wilson et al., 2016). Therefore, it is reasonable to assume that in mammalian *PTPRJ* the translating ribosomes stall when the arginine residues of the nascent chain are in the exit tunnel. The result is translation inhibition. The high degree of conservation of this inhibitory sequence is a strong indication of its functional significance.

PTPRJ has numerous cellular substrates, such as RTKs, Src-family kinases, and others. It has been implicated in the regulation of a wide range of cellular functions. Enzymatic studies of the PTPRJ phosphatase domain revealed promiscuity with respect to substrate specificity and a very high intrinsic activity (Barr, 2009). It appears therefore plausible that high levels of PTPRJ protein may disturb cellular functions or may even be toxic. Attenuation of PTPRJ translation by virtue of the here described features of the signal peptide may serve to prevent toxic effects and to allow fine-tuning of expression at the transcriptional level.

Other transcripts, encoding N-terminally extended signal peptides, may use a similar cis-acting mechanism to attenuate expression. An analysis of the extended signal peptides in the precursors of the human E3 ubiquitin-protein ligase ZNRF3 and NOTCH3 revealed the presence of arginine clusters and stretches of proline residues. Both structures may cause ribosome pausing as the nascent chain is synthesized (Fig. 4). In these cases, the efficiency of the nascent chains to throttle translation remains to be elucidated.

**Figure 4.**
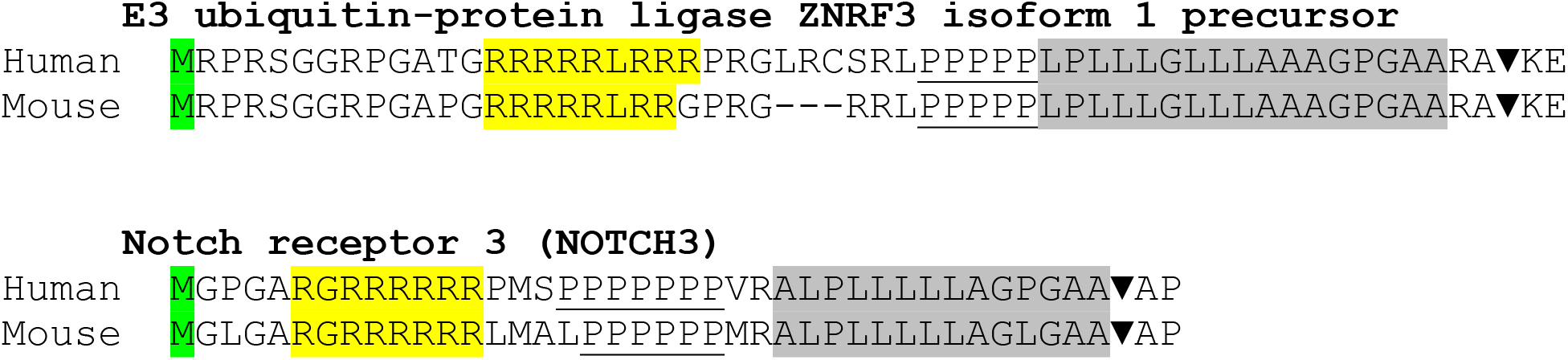
The N-terminally extended signal peptide of the E3 ubiquitin-protein ligase ZNRF3 and NOTCH3 in humans and mouse. The initiating Met residues (green), the Arg-cluster (yellow), the consecutive Pro residues (underlined) and the nonpolar signal sequences (grey) are shown. The signal peptidase cleavage site (▼) was predicted *in silico* by the SignalP 5.0 Server.

## Acknowledgments

One of the authors (LK) highly appreciates the help of I. Stancheva for critically reading the manuscript and helpful suggestions.

