## Supplemental Figures for "The translation attenuating arginine-rich sequence in the extended signal peptide of the protein-tyrosine phosphatase PTPRJ/DEP1 is conserved in mammals"

### Supplementary figures

Supplementary Fig. 1.

Alignment of the 5′ end of the *PTPRJ* mRNAs encoding the extended signal peptides in placental mammals.

The AUG/ATG triplets (green) upstream of the hydrophobic signal sequence are in-frame with the main protein. In the consensus sequence, the AUG/ATG codons and the flanking nucleotides are underlined. The regions encoding the conserved Arg-cluster (yellow) and the nonpolar signal sequence (grey) are marked. The signal peptidase cleavage sites were predicted *in silico* by the SignalP 5.0 Server. The end of exon 1 in human *PTPRJ* is marked (▼).

```
Human          1 ---AGCCCCAGCCGCATGACGCGCGGAGGAGGCAGCGGGA
Mouse          1 -------GGAGCCGCATGACGCGCGGAGGAGGCAGAGGGA
Beluga         1 TGCAGCCCCATCCGCATGACGCGCGGAGGCGGCAGCCGGA
Cattle         1 ---------AGCCGCATGACGCGCGGAGGAGGCAGCGGGA
Otter            ----------------------------------------
Consensus                 AGCCGCATGACGCGCGGAGGAGGCAGCGGGA

Human         38 GCAGCCGCGGG------AGCCGGGACCGGGTAGCCGCGCG
Mouse         34 GCAGCCGGGGCCGCGGGAGCCGGGAGCTGGGAGCCACGCG
Beluga        41 GCCGCCGG---------AGCCGGGATCAGGGCGCTGAGCG
Cattle        32 GCCGCCGGGGCCGCGGGAGCCGGGATCCGGGCGCCGCGCG
Otter            ----------------------------------------
Consensus        GC GCCGGGG       AGCCGGGA C GGG GCCGCGCG

Human         72 CTGGGGGTGGGCGCCGCTCGCTCCGCCCCGCGAAGCCCCT
Mouse         74 CGGAGGGTGGGCGCCGCTCGCTCCGCCCCGCGAAGCCCCT
Beluga        72 CGGGGGGTGGGCGCCGCTCGCTCCGCCCTGCGAAGCCCCT
Cattle        72 CCAGGGGTGGGCGCCGCTTGCTCCGCCCTGCGAAGCCCCT
Otter          1 --------------------------------AAGCCCCT
Consensus        C GGGGGTGGGCGCCGCTCGCTCCGCCC GCGAAGCCCCT

Human        112 GCGCGCTCAGGGACGCGGCCCCCCCGCGGCAGCCGCGCTA
Mouse        114 GCGAGTCTAAGGCCGCGGCCGCTCCGCGCCAGGCGCGCTA
Beluga       112 GCGCGCCCAGGGACGCGTCCACCCCGCTGCAGCCCCGCCA
Cattle       112 AGGCGTCCGGGGACGCGTTCCCCCCGCCGCAGCCGCGCCG
Otter          9 GCGCGCCCAGGGACGCGTCCCTCCCGCGGCAGGCGCGCCA
Consensus        GCGCGCCCAGGGACGCGTCCCCCCCGCGGCAGCCGCGCCA

Human        152 GGCTCCGGCGTGTGGCCGCGGCCGCCGCCGCCGC---TGC
Mouse        154 GGCTCCGCCGTGTG------GCCGCCGCCGCCGCCGCTGC
Beluga       152 GGCTCCGCCGTGTGGCCGCTGCCGCCGCCGCTGC---TGC
Cattle       152 GGCTCCGCCGTGTG------GCCGCCGCCGCCGCCGCTGC
Otter         49 GGCTCCGCCGTGTGGCCGCCGCCGCCGCCGCCGCCGCTGC
Consensus        GGCTCCGCCGTGTGGCCGC GCCGCCGCCGCCGCCGCTGC

Human        189 CATGTCTCCGGGGAAGCCCGGGGCGGGCGGAGCGGGGACG
Mouse        188 CATGTCCCCGGGGAAGCCCGGGGCGGGCGGAGCGGGGACT
Beluga       189 CATGTCCCCGGGGAAGCCCGGGGCGGGCGGAGCGGGGACG
Cattle       186 CATGTCCCCGGGGAAGCCCGGGGCGGGCGGAGCGGGGACG
Otter         89 CATGTCCCCGGGGAAGCCCGGGGCGGGCGGAGCGGGGACT
Consensus        CATGTCCCCGGGGAAGCCCGGGGCGGGCGGAGCGGGGACG

Human        229 AGGCGGACCGGCTGGCGGAGGAGGAGGCGAAGGAGACGGC
Mouse        228 AGGCGGACCGGCTGGCGGAGAAGGAGGCGGAGGCGTCGGC
Beluga       229 AGGCGGACCGGCTGGAGGAGGAGGAGGCGGAGGCGGCGGC
Cattle       226 AGGCGGACCGGCGGGAGGAGGCGGAGGCGGAGGCGGCGGC
```

```
Otter          129 AGGCGGACGGGCTGGAGGAGGAGGAGGAGGAGGCGGCGGC
Consensus          AGGCGGACCGGCTGGAGGAGGAGGAGGCGGAGGCGGCGGC

Human          269 AGGAGGCGGCGACGACGGTGCCCGGGCTCGGGCGCACGGC
Mouse          268 TGGAGACGGAGACGAGGGCGCCCGGCTTCGGGCACACGGC
Beluga         269 TGGAGGCGGCGACTAGGGCGCCGGGGCTCGGAAGAACGGT
Cattle         266 TGGAGGCGGTGACTCAGGCGCCGGGGCTCGGGCGCACGGC
Otter          169 TAGAGGCGGCGACTAGGGCGCCCGGGGTCGGGCGCACGGC
Consensus          TGGAGGCGGCGACTAGGGCGCCCGGGCTCGGGCGCACGGC

Human          309 GGGGCCCGATTCGCGCGTCCGGGGCACGTTCCAGGGCGCG
Mouse          308 GGG---------GCGCGTCCCGGGCACGTTCCAGGGCGCG
Beluga         309 GGGGCCCTACTCGCGCGTCCCGGGCACGTTCCAGGGCGCG
Cattle         306 GGGGCCCGGCTCGCGCGTCCCGGGCACGTTCCAGGGCGCG
Otter          209 GGGGCCCGACTCGCGCGTCCCGGGCACGTTCCAGGGCGCG
Consensus          GGGGCCCGACTCGCGCGTCCCGGGCACGTTCCAGGGCGCG

Human          349 CGGGGCATGAAGCCGGCGGCGCGGGAGGCGCGGCTGCCTC
Mouse          339 CAGGGCATGAAGCCCGCGGCGCGGGAGACGCGGACACCCC
Beluga         349 CGGGGCATGAAGCAGGCGGCGCGGGAGGCGCGGCCGCCTC
Cattle         346 CGGGGCATGAAGCCGGCGGCTCGGGAGGCGCGGCCGCCTC
Otter          249 CGGGGCATGAAGCGGGCGACGCGGGAGGCGCGGCCGCCTC
Consensus          CGGGGCATGAAGCCGGCGGCGCGGGAGGCGCGGCCGCCTC

Human          389 CGCGCTCGCCCGGGCTGCGCTGGGCGCTGCCGCTGCTGCT
Mouse          379 CGCGCTCGCCCGGGCTCCGCTGGGCGCTGCTGCCGCTGCT
Beluga         389 CGCGCTCGCCCCGGCTCCGCTGGGCGCTGCCGCCGCTGCT
Cattle         386 CGCGCTCGCCCGGGCTGCGCTGGGCGCTGCCGCCGCTGCT
Otter          289 CGCGCTCGCCGGGGCTGCGCTGGGCGCTGCCGCCGCTGCT
Consensus          CGCGCTCGCCCGGGCTGCGCTGGGCGCTGCCGCCGCTGCT
                             first exon ▼ second exon
Human          429 GCTGCTGCTGCGCCTGGGCCAGATCCTGTGC------
Mouse          419 GCTGTTGCTACGCCAGGGCCAGGTCCTGTGC------
Beluga         429 GCTGCTGTTGCGCCTGGGCCAGATCTTGTGTACAGGT
Cattle         426 GCTGCTGTTGCGCTTGGGCCAGATCCTGTGC------
Otter          329 GCTGCTGTTGCGCCTGGGCCAGATGGTGTGCGCA---
Consensus          GCTGCTGTTGCGCCTGGGCCAGATCCTGTGC
```

Supplementary Fig. 2.

Alignment of the PTPRJ N-terminally extended signal peptides in placental mammals. Translation starts at the second AUG triplet in the mRNA (see Suppl. Fig. 1). The initiating Met residues (green), the conserved Arg-clusters (yellow) and the nonpolar signal sequences (grey) are marked. The cleavage sites (▼) of the signal peptidase were predicted by the SignalP 5.0 Server *in silico*.

```
Human          1  MSPGKPGAGGAGTRRTGWRRRRRRRRQEAATTVPGLGRTA
Mouse          1  MSPGKPGAGGAGTRRTGWRRRRRRRRLETETRAPGFGHTA
Beluga         1  MSPGKPGAGGAGTRRTGWRRRRRRRRLEAATRAPGLGRTV
Cattle         1  MSPGKPGAGGAGTRRTGGRRRRRRRRLEAVTQAPGLGRTA
Otter          1  MSPGKPGAGGAGTRRTGWRRRRRRRRLEAATRAPGVGRTA
Consensus         MSPGKPGAGGAGTRRTGWRRRRRRRRLEAATRAPGLGRTA

Human         41  GPDSRVRGTFQGARGMKPAAREARLPPRSPGLRWALPLLL
Mouse         41  G---RVPGTFQGAQGMKPAARETRTPPRSPGLRWALLPLL
Beluga        41  GPYSRVPGTFQGARGMKQAAREARPPPRSPRLRWALPPLL
Cattle        41  GPGSRVPGTFQGARGMKPAAREARPPPRSPGLRWALPPLL
Otter         41  GPDSRVPGTFQGARGMKRATREARPPPRSPGLRWALPPLL
Consensus         GP SRVPGTFQGARGMKPAAREARPPPRSPGLRWALPPLL

Human         81  LLLRLGQILC▼AG
Mouse         78  LLLRQGQVLC▼AG
Beluga        81  LLLRLGQILCTG▼DC
Cattle        81  LLLRLGQILC▼AD
Otter         81  LLLRLGQMVCA▼GD
Consensus         LLLRLGQILC
```

Supplementary Fig. 3.

Alignment of the 5' end of the *PTPRJ* transcripts encoding the extended signal peptides in marsupials. The AUG/ATG codons (green), the regions encoding the conserved Arg-cluster (yellow) and the nonpolar signal sequence (grey) are marked. In the consensus sequence, the AUG/ATG codons and the flanking nucleotides are underlined. The signal peptidase cleavage sites were predicted *in silico* by the SignalP 5.0 Server.

```
Opossum          1 AGGGAGGGGGCTGGCTTCTCCCGGAGGTGGCGGCTGC---
Koala            1 -GGGAGGGGGTTGGCTTCTCCC---GGCGGCTGCTGC---
T. devil         1 ---GGGGGGGCTGGCTTCTCCCGGCGGCGGCGGCGGCGGC
Wombat             ----------------------------------------
Consensus           ggGaGGGGGcTGGCTTCTCCCgg GG GGCgGCtGC

Opossum         38 AGGGAGCCCGAGCAGCGGGAGCCGCCGGAGCCC---GAG-
Koala           34 AGGGAGCCCGAGCCACGGGAGCCGCCGGAGCCA---CCGG
T. devil        38 AGGGAGCCCGAGCCGCGGGAGCCTCCGGAGCCACCGGAG-
Wombat             ----------------------------------------
Consensus          AGGGAGCCCGAGC gCGGGAGCCgCCGGAGCC    gaG

Opossum         74 -----CCCGAGCCCGAGC----A-CCGG--------AGCC
Koala           71 AGCCGCCGGAACCCGAGC----AGCCGG--------AGCC
T. devil        77 -----CCCGAGCGCGAGCCGGAA-CCGAGCAGTCGCAGCC
Wombat             ----------------------------------------
Consensus               CCcGAgCcCGAGC    A CCGg        AGCC

Opossum         96 CAAGCCGCAGGACGCGTGGAGTAGGCAGCGGGAGCCCGAG
Koala           99 CGAGCCGCAGGACGCGTGGAGCAGGCAGCTGGA-------
T. devil       111 CGAACCACAGGAGGCGTGGAGTCGGCAGCTGGAGCCCGAG
Wombat             ----------------------------------------
Consensus          C AgCCgCAGGAcGCGTGGAGtaGGCAGC GGAgcccgag

Opossum        136 CCGCCCGAGCCGCCCAAGCCGCCCTAGCCTGGACGCCCCC
Koala          132 -----------GCCCAAGCCGCCCGAGCCTGGACGCCGCC
T. devil       151 CCGCCCG-------------------AGCCTG--------
Wombat             ----------------------------------------
Consensus          ccgcccg    gcccaagccgccc AGCCTGgacgcc cc

Opossum        176 GCCCCCGCCGCCGCCGCGCCTCTGCAGCTCGGGGGGGTGG
Koala          161 GCCTCCGCCGCCGCCGCGCCGCTGCAGCTC-GGGGGGTGG
T. devil       164 ------GCCGCCGCCGCGCCGCTGCAGCTCG-GGGGGTGG
Wombat             ----------------------------------------
Consensus          gcc ccGCCGCCGCCGCGCC CTGCAGCTCggGGGGGTGG

Opossum        216 GCGCCGCTTGCTCCGCCCCGTGGAAGCCCTCCTGGCCGCC
Koala          200 GCGCCGCTCGCTCCGCCCCGTGGAAGCCCTCCTGGCCGCC
T. devil       197 GCGCCGCTCGCTCCGCCCCGTGGAAGCCCTCCCGGCCGCC
Wombat             ----------------------------------------
Consensus          GCGCCGCT GCTCCGCCCCGTGGAAGCCCTCCtGGCCGCC
```

```
Opossum        256 GCCACCCCCGCCGCCTTCTTCACCGAGGGACTCGGGGCTG
Koala          240 GCCGCCTCCG------------CCGAGGGACGCGGGGCTG
T. devil       237 GCCGCCGCCTCCG---------CCGAGGGACGCGGGGCTG
Wombat             ----------------------------------------
Consensus          GCC CC CCgccg         CCGAGGGAC CGGGGCTG

Opossum        296 CTCCCGCCCGTTCGAGACTCAGCCTGGTGGTGGCCGCCGC
Koala          268 CTCCCGCCCGCCCGAGACCCAGCCTGGTGGTGGCCGCCGC
T. devil       268 CTCCCGCCCGCCCGAGACCCAGGCTGCCGGTGGCCGCCGC
Wombat             ----------------------------------------
Consensus          CTCCCGCCCG  CGAGAC CAGcCTGgtGGTGGCCGCCGC

Opossum        336 ---TGCCATGTCCCCGGGGAAGCCCGGGGCGGGCGGAGCG
Koala          308 CGCTGCCATGTCCCCGGGGAAGCCCGGGGCGGGCGGAGCG
T. devil       308 CGCTGCCATGTCCCCGGGGAAGCCCGGGGCGGGCGGAGCG
Wombat           1 -------ATGTCCCCGGGGAAGCCCGGGGCGGGCGGAGCG
Consensus             TGCCATGTCCCCGGGGAAGCCCGGGGCGGGCGGAGCG

Opossum        373 GAGAGGAGGCGGAGGAGCTGGAGGCGGCGGCGGCGGCGGC
Koala          348 GAGAGGAGGCGGAGGAGCTGGAGGCGGCGTCGGCGGCGGC
T. devil       348 GAGAAGAGGCGGAGGACGTGGAGGCGGCGGCGGCGGCGGC
Wombat          34 GAGAGGAGGCGGAGGAGCTGGAGGCGGCGGCGGCGGCGGC
Consensus          GAGAGGAGGCGGAGGAGCTGGAGGCGGCGGCGGCGGCGGC

Opossum        413 GGCCTCGGCCCCGGCCACCGGCTCCAGCT-----------
Koala          388 GGCCTCGGCCCCGGCCACCGGCTC----------------
T. devil       388 GGCCTCGGCCCGGGCCCCCGGCTGCGGCTCCCGGAGCCCA
Wombat          74 GGCCTCGGCCCCGGCCACCG------GCT-----------
Consensus          GGCCTCGGCCCCGGCCACCGGCTcc GCT

Opossum        442 -------GCTGTCCCCGGCGCCGAGGCTGCTGCAG---AA
Koala          412 --------CTGCCCCGGGCGCCGAGGCTGCTGCTG---AA
T. devil       428 GGCTGCTGCGGCTCCCGGAGCCCAGGCTGCTGCCGCTGAA
Wombat          97 -------GCTGCCCCGGGCGCCGAGGCTGCTGCTG---AA
Consensus                 GCTG CCCcGGCGCCGAGGCTGCTGC G   AA

Opossum        472 GCTGCTGCTCTGGGGCCCCAGAGAGCCGCGCTGCTCCCGG
Koala          441 GCTGCTGCTCCGGGGCCCCGGCGCGCCGCGCTGCTCCCGG
T. devil       468 GCCGCCGCTCCGGGGCCCCGCCGCGCCGCGCCGCTCCCGG
Wombat         127 GCTGCTGCTCCGGGGCCCCGGCGCGCCGCGCTGCTCCCGG
Consensus          GCTGCTGCTC GGGGCCCC G G GCCGCGCTGCTCCCGG

Opossum        512 GCACGTTCAGGGACGCTCGGAGCCCGAAGCCCGGGGGGGC
Koala          481 GCTCGTTCCGGGGCGCGCGGAGCCCGAAGCCCGGGGGGGC
T. devil       508 GCTGGTTCCGGGGCGCGCGGAGCCCCAAGCTCGGGGAGGC
Wombat         167 GCTCGTTCCGGGGCGCGCGGAGCCCGAAGCCCGGGTGGGC
Consensus          GC CGTTC GGG CGC CGGAGCCCGAAGCCCGGGGGGGC

Opossum        552 GGGGGCGAAGCCTCCGCTCTGTCTCCTGCTGCGCCTGGGG
Koala          521 GGGGGCGCAGCCTCGGCTCTGCCTCCTGCTGAGCCTGGGG
T. devil       548 GGGGGCGCAGCCTCGGCTCTGTCTCCTCCTGCGCCTGGGG
Wombat         207 TGGGGCGCAGCCTCGGCTCTGCCTCCTGCTGCGCCTGGGG
```

```
Consensus           GGGGGCG AGCCTC GCTCTGtCTCCTGCTGCGCCTGGGG

Opossum        592 CTGCTATTGCTCCGCTTCAGCCAGATTGCTGTTGCA
Koala          561 TTGCTACTGCTCCGCTTCGGCCAGATTGCAGTTGCA
T. devil       588 CTGCTCCTGCTCGCCTTCTGCCAGATTGCTGTTGCA
Wombat         247 CTGCTACTGCTCGGCTTCGGCCAGATTGCAGTTGCA
Consensus          CTGCTA TGCTCcGCTTC GCCAGATTGCtGTTGCA
```

Supplementary Fig. 4.

Alignment of the N-terminally extended signal peptides in PTPRJ of marsupials.
The initiating Met residues (green), the conserved Arg-clusters (yellow) and the nonpolar signal sequences (grey) are marked. The cleavage sites (▼) of the signal peptidase were predicted *in silico* by the SignalP 5.0 Server.

```
Opossum          1 MSPGKPGAGGAERRRRSWRRRRRRRPRPRPPAPA------
Koala            1 MSPGKPGAGGAERRRRSWRRRRRRRPRPRPPAPA------
T.devil          1 MSPGKPGAGGAEKRRRTWRRRRRRRPRPGPPAAAPGAQAA
Wombat           1 MSPGKPGAGGAERRRRSWRRRRRRRPRPRPPA--------
Consensus          MSPGKPGAGGAERRRRSWRRRRRRRPRPRPPApA

Opossum         35 AVPGAEAAA-EAAALGPQRAALLPGTFRDARSPKPGGAGA
Koala           35 --PGAEAAA-EAAAPGPRRAALLPGSFRGARSPKPGGAGA
T.devil         41 AAPGAQAAAAEAAAPGPRRAAPLPGWFRGARSPKLGEAGA
Wombat          33 AAPGAEAAA-EAAAPGPRRAALLPGSFRGARSPKPGWAGA
Consensus          A PGAEAAA EAAA GP RAALLPG FR ARSPKPGgAGA

Opossum         74 KPPLCLLLRLGLLLLRFSQIAVA▼VS
Koala           72 QPRLCLLLSLGLLLLRFGQIAVA▼GD
T.devil         81 QPRLCLLLRLGLLLLAFCQIAVA▼DD
Wombat          72 QPRLCLLLRLGLLLLGFGQIAVA▼GD
Consensus           P LCLLLRLGLLLLrF QIAVA
```

Supplementary Fig. 5.

The 5’ end region of the *PTPRJ* mRNA in platypus (monotremes) encoding the extended signal peptide. The in-frame AUGs triplets upstream of the nonpolar signal sequence (green), the region encoding the Arg-cluster (yellow) and the nonpolar amino acid signal sequence (grey) are marked. In the sequence, the AUG/ATG codons and the flanking nucleotides are underlined. The cleavage site of the signal peptide was predicted *in silico* by the SignalP 5.0 Server.

```
  1   GGCTGCAAAA GCAGCAGGAG CTGCAGTTGC AGCAATCGCA GCAGCAGCAG
 51   CAATTGCAAC AGCAGTTGCA GCAGCAGCAG CAATTGCAAC CTCCGCAGCG
101   GCAATTGCAG CAGCAACGGC CGGGCCGCGG TCCCTCCCCC CCGCGCCAAC
151   CCTGCCCGTT GCGGCCGGCC GAGGGCTGCT GCGGGGCCGT TGCGGCCGGA
201   GGCGCGCCCG CCCCCCCGCC ATGTCCCCGG GGAAGCCCGG AGCGGGGGAG
251   ACGCCTCCGA GGAGGAGGAG GCGGCGGGGG AGGCGGAGGA GGAGGAGGAG
301   GAGGCCCCAG CCGGGACCGG CGACGACGAA GGCGGCGGCG GGTGGAGCCG
351   GGCCCCGGCT TGCGGGCCTC CCGGGAAGGC TGGGCGGCAT GAAGCTCGGC
401   TCCCTGCTGG GGCTGCTCTT GCTGCTGCAT TCCGGACAGA TGAGATGT
```

Supplementary Fig. 6.

The N-terminally extended signal peptide of *PTPRJ* in platypus (monotremes).
The initiating Met residue (green), the Arg-cluster (yellow), and the nonpolar signal sequence (grey) are shown. The signal peptidase cleavage site (▼) was predicted *in silico* by the SignalP 5.0 Server.

```
 1   MSPGKPGAGE TPPRRRRRRG RRRRRRRRPQ PGPATTKAAA GGAGPRLAGL
51   PGRLGGMKLG SLLGLLLLLH SGQMRC▼AG
```
